# Boldine inhibits SARM1 NADase Activity and Preserves Axonal Integrity After Nerve Injury

**DOI:** 10.64898/2026.07.27.741059

**Authors:** Justin C. Burrell, Tomisin T. Latona, Henry Garcia, David R. Clizbe, Mykhailo M. Tatarchuk, Liwen Zhou, Carlos A. Toro, Alexa N. Olivarez, Phuong V. Nguyen, Cathy Z. Yang, George D. Bittner, Christopher P. Cardozo, D. Kacy Cullen

## Abstract

Wallerian degeneration of anucleated axonal segments is driven by SARM1, which depletes axonal NAD+, disrupting energy metabolism and triggering self-destruction. SARM1 inhibition is an emerging therapeutic target for traumatic nerve injuries. Boldine, a natural aporphine alkaloid from *Peumus boldus*, modulates connexin hemichannels, oxidative stress, and inflammation. Building on our published work showing boldine’s neuroprotective effects in nerve injury models, we hypothesized that boldine also inhibits SARM1 directly. A fluorescence polarization assay revealed that boldine inhibits SARM1 NADase activity with an IC50 of approximately 7.5 uM. AI-assisted structural modeling (AlphaFold3-based Boltz-1 with GNINA docking) predicted two boldine binding sites on SARM1: the TIR catalytic site (Kd ∼ 13.5 uM) and the ARM-TIR regulatory interface (Kd ∼ 12 uM). In a sciatic nerve explant model, boldine preserved the integrity of anucleated axonal segments at 3 and 7 days post-transection relative to vehicle controls. These findings suggest boldine may act as a dual-site SARM1 inhibitor and support its development as a neuroprotective therapy after traumatic axonal injury.

## Introduction

Traumatic nerve injury initiates a rapid cascade of degeneration of anucleated axonal segments that precedes and often dictates long-term functional outcomes.^1, 2^ In both peripheral and central nervous system injuries, disruption of axonal integrity leads to loss of electrical conduction, denervation of target tissues, and progressive functional decline, even when neuronal cell bodies remain viable. Rather than a passive consequence of damage, axonal degeneration is now understood as an active, tightly regulated self-destruction program that is triggered within hours of injury and propagates along the distal axonal segment.^3–5^ This early degenerative phase represents a critical therapeutic window, as preservation of axonal structure and function has the potential to maintain connectivity, prolong tissue viability, and improve the likelihood of successful regeneration and reinnervation.^2^

At the center of this degenerative program is SARM1 (Sterile Alpha and TIR Motif Containing 1), a multidomain NAD⁺ hydrolase that functions as a metabolic checkpoint for axonal survival.^6, 7^ Genetic deletion of SARM1 in rodent models markedly delays Wallerian degeneration, preserving axonal integrity for weeks after injury and establishing SARM1 as a central driver of injury-induced axonal loss.^8–12^ These findings have positioned SARM1 as a compelling therapeutic target for neuroprotection across traumatic nerve injuries, peripheral neuropathies, and neurodegenerative conditions.

SARM1 comprises three key domains: an N-terminal Armadillo repeat motif (ARM) domain that regulates autoinhibition and senses metabolic state, tandem sterile alpha motif (SAM) domains that promote oligomerization, and a C-terminal Toll/Interleukin-1 receptor (TIR) domain that contains the NADase catalytic site.^13–15^ Under basal conditions, the ARM domain suppresses TIR activity, maintaining SARM1 in an inactive conformation. Following injury, rapid loss of NMNAT2 leads to a shift in the intracellular NMN:NAD⁺ ratio, which relieves ARM-mediated autoinhibition and promotes SAM-dependent oligomerization.^10, 16–18^ Notably, human SARM1 assembles as a homo-octamer, and the multimeric architecture is central to both its autoinhibited resting state and its activation-competent conformation. This conformational transition activates TIR NADase activity, resulting in rapid NAD⁺ depletion and bioenergetic collapse, culminating in axonal fragmentation typically within 24–48 hours in experimental models.^15, 19–21^

Three principal pharmacological strategies have emerged to inhibit SARM1, each with distinct limitations.^22^ Covalent allosteric inhibitors targeting an ARM-domain cysteine (Cys311), such as tryptoline acrylamides, lock SARM1 in its autoinhibited conformation and show high proteome-wide selectivity; however, concerns about irreversibility and potential off-target reactivity remain.^23, 24^ Covalent orthosteric inhibitors targeting the TIR NADase domain, including electrophilic isothiazoles that modify catalytic cysteine residues, have demonstrated robust axon-protective effects in preclinical models, but raise concerns regarding off-target reactivity, immunogenicity, and the inherent risks associated with irreversible enzyme modification.^23, 25^ Base-exchange modulators exploit SARM1’s catalytic mechanism to generate inhibitory ADPR adducts in situ, yet paradoxically can activate the enzyme at sub-inhibitory concentrations, potentially narrowing the therapeutic window.^26^ Collectively, these challenges underscore the need for reversible inhibitors based on well-characterized scaffolds that can engage SARM1 while maintaining favorable pharmacological properties. Notably, the recent advancement of a SARM1 inhibitor into Phase I clinical trials highlights the translational momentum in this area.^22^

Boldine ([S]-2,9-dihydroxy-1,10-dimethoxyaporphine), an aporphine alkaloid derived from the Chilean boldo tree (*Peumus boldus*), represents a structurally distinct candidate for SARM1 inhibition with an established safety profile and demonstrated central nervous system effects, including blood–brain barrier penetration in rodent models.^6, 27, 28^ Beyond its antioxidant and anti-inflammatory activities, boldine blocks movement of small molecules through open connexin hemichannels and has shown neuroprotective effects across models of ischemia, neuroinflammation, and excitotoxic injury (**Fig. 1**).^29–32^ The lipophilic, planar aporphine scaffold of boldine is well suited for engagement with hydrophobic and interfacial binding pockets, supporting its evaluation as a potential SARM1 ligand.^33, 34^

**Fig 1.**
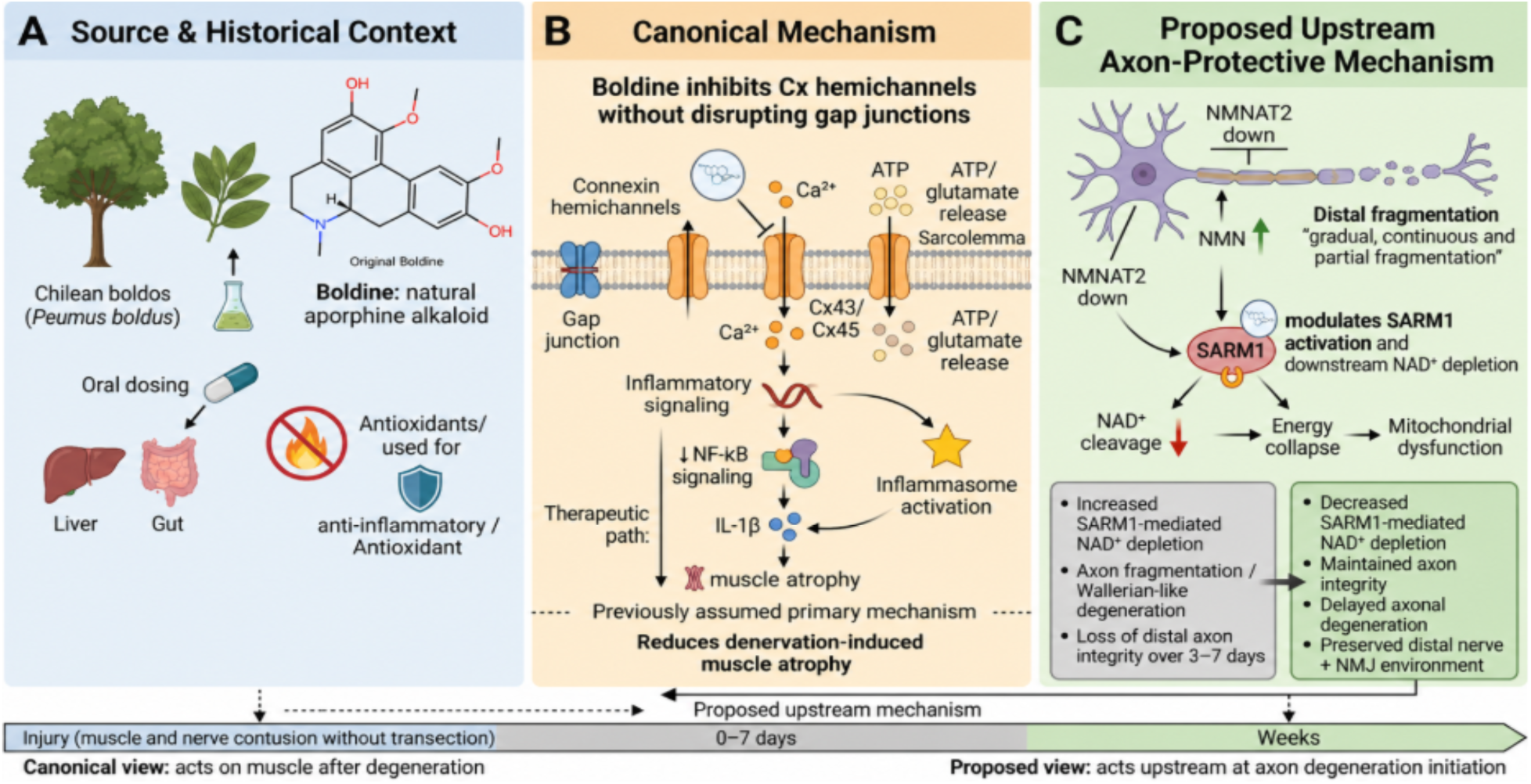
Conceptual framework for boldine mechanism of action in neuromuscular injury. (A) Source and historical context. Boldine is a naturally occurring aporphine alkaloid derived from the Chilean boldo tree (Peumus boldus), traditionally used for its antioxidant and anti-inflammatory properties following oral administration. (B) Canonical mechanism. Boldine has been shown to inhibit connexin (Cx) hemichannels (Cx43/Cx45) in the sarcolemma, reducing calcium influx and ATP release. This attenuates downstream inflammatory signaling, including NF-κB activation, inflammasome activity, and IL-1β production, thereby decreasing denervation-induced muscle atrophy. Note: skeletal muscle fibers do not express functional gap junctions or glutamate receptors; the canonical boldine target in this context is the Cx43/Cx45 hemichannel, not gap junctions or glutamate signaling. ^50^ (C) Proposed mechanism. Following nerve injury, depletion of NMNAT2 and accumulation of NMN activate SARM1, leading to NAD⁺ cleavage, mitochondrial dysfunction, and axonal degeneration. In the absence of treatment, SARM1 activation drives rapid distal axon fragmentation and loss of neuromuscular integrity. With boldine treatment, SARM1-mediated NADase activity is reduced, preserving axonal structure, delaying degeneration, and maintaining the distal nerve and neuromuscular junction (NMJ) environment. Bottom timeline: The canonical view places boldine action downstream at the level of denervated muscle following degeneration, whereas the proposed mechanism suggests an upstream effect at the initiation of axonal degeneration, shifting the therapeutic target from muscle preservation to axon protection.

We hypothesized that simultaneous engagement of the TIR NADase catalytic site and the ARM–TIR regulatory interface could provide mechanistic redundancy across orthosteric and allosteric axes. This hypothesis was evaluated through integrated computational modeling, using AI-enabled structural biology approaches that combined AlphaFold3-based modeling (via Boltz-1) with deep learning–guided molecular docking (GNINA) to predict potential boldine–SARM1 interactions, together with biochemical NADase activity assays using fluorescence polarization and morphological assessment in an *ex vivo* nerve injury model. While the results are consistent with boldine acting as a naturally occurring dual-site SARM1 inhibitor, further direct structural validation will be required to definitively establish this boldine mechanism of action.

## Methods

### SARM1 NADase Activity Assay

#### Assay principle

SARM1 NAD+ hydrolase activity was measured using the far-red competitive fluorescence polarization (FP) immunoassay kit, Transcreener ADPR FP (BellBrook Labs, Cat #3030-1K). In this method, SARM1 cleaves NAD^+^ to produce ADPR; this ADPR is then converted in real time to AMP by an ADPR-AMP coupling enzyme. The generated AMP competes with an AMP2/GMP2 fluorescent tracer for binding to an AMP2/GMP2 antibody, resulting in a detectable change in FP signal. The assay is designed for biochemical measurements with purified enzyme.

#### Reagents and instrumentation

Recombinant human SARM1 (BPS Bioscience, Cat #100069) was used with an assay buffer containing 50 mM Tris-HCl (pH 7.5), 5 mM MgCl₂, and 0.01% Triton X-100, prepared in nuclease-free water. Reactions were performed in black 384-well plates and measured on an FP-capable plate reader (iD5 Molecular Devices).

#### Reaction setup

Enzyme reactions were prepared in a total volume of 10 μL (consisting of 5 μL enzyme/buffer and 5 μL substrate/coupling mix) and then expanded to a 20 μL “complete assay” by adding 10 μL of 1X AMP Detection Mix. NAD^+^ was used at a final concentration of 20 μM (from a 40 μM working concentration in the substrate/coupling mix), with coupling enzyme at 1X in the enzyme reaction. The detection mix resulted in final “complete assay” concentrations of 0.5X Stop & Detect Buffer B, 4 nM AMP2/GMP2 tracer, and 7.5 μg/mL AMP2/GMP2 antibody. Plates were mixed (∼40 s), sealed, incubated for 60 minutes at 30°C for the enzyme reaction, then incubated for 90 minutes at room temperature after adding the detection mix before measuring FP.

#### Enzyme titration, ADPR standard curve, and boldine inhibitor dose response

To determine the appropriate enzyme concentration, SARM1 was titrated using two-fold serial dilutions across wells, aiming to stay within the initial-velocity regime (<20% substrate conversion) and selecting a concentration that yielded approximately 80% of the maximal signal. For quantitative conversion of the FP signal to the product formed, an 11-point ADPR standard curve was created by serially diluting ADPR (starting from a 40 μM working solution) to cover a range from 20 μM down to 0 μM. For inhibitor profiling, SARM1 was preincubated with test compounds (30 min, room temperature) before initiating reactions with substrate/coupling mix and proceeding as described above; IC₅₀ values were determined from dose-response curves.

### Protein Structure Preparation

Protein structures for SARM1 were obtained from the RCSB Protein Data Bank (PDB) and included the inactive octameric complex (PDB ID: 7CM6), the activated full-length enzyme (PDB ID: 7NAL), the isolated ARM regulatory domain (PDB ID: 7M6K), and the TIR catalytic domain (PDB ID: 6O0Q).^7, 9, 13, 14, 16, 19, 35, 60^ Additionally, structural ensembles were generated using the AlphaFold3-based Boltz-1 model accessed via the Neurosnap computational platform.^34, 36^ Protein structures were prepared for docking by removing water molecules, adding hydrogen atoms, and assigning Gasteiger partial charges using UCSF Chimera.^37^ Missing loops and side chains were modeled using MODELLER.^38^

### Ligand Preparation

The three-dimensional structure of boldine was obtained from PubChem (CID: 10154) and energy-minimized using the MMFF94 force field in Avogadro.^39^ Rotatable bonds were identified, and torsional degrees of freedom were assigned. Known SARM1 inhibitors that served as control compounds (dehydronitrosonisoldipine^63^ and DSRM-3716^18, 19^) were prepared following the same protocol.

### Molecular Docking Protocol

Molecular docking was performed using GNINA version 1.3, a deep learning framework that employs 3D convolutional neural networks (CNNs) to score protein-ligand interactions.^33, 40^ We set an exhaustiveness level of 8 and generated 10 binding poses for each ligand. For blind docking experiments on the TIR domain, the entire surface of the domain was included in the search space. For targeted docking at specific pockets, the search box was centered on key residues identified through structural analysis (e.g., Glu642, His685, Asn640 for the NAD⁺ binding region).

### Scoring and Thermodynamic Calculations

Binding affinities were predicted using the CNN scoring function in GNINA.^33, 40^ Thermodynamic parameters were calculated with the equations: ΔG = −RT ln(K*a*); K*d* = 1/K*a*, where R is the gas constant (1.987 cal·mol⁻¹·K⁻¹), T is the temperature (298 K), and K*a* is the association constant. The top-scoring pose for each ligand was selected for detailed structural analysis and visualization in UCSF Chimera and PyMOL.^37^

### Validation of Docking Protocol

The docking protocol was validated by re-docking known SARM1 inhibitors (DSRM-3716) and comparing predicted K*d* values with experimentally determined binding constants reported in the literature.^41^ Validation was performed using the TIR domain structure (PDB ID: 6O0Q) containing a co-crystallized ligand, with the same search box and exhaustiveness settings used for boldine docking. Re-docking was performed without prior knowledge of the crystallographic pose (i.e., the pose was not used to center the search box, which was instead defined by the catalytic residues Glu642, His685, and Asn640). Protonation states were assigned at physiological pH (7.4) using Gasteiger charges in UCSF Chimera before docking.

The root-mean-square deviation (RMSD) between predicted and crystallographic poses was less than 2.0 Å, supporting the reliability of the method under these conditions.

### *Ex vivo* Nerve Culture and Axon Degeneration Assay

#### Ethics statement

All animal procedures were approved by the Corporal Michael J. Crescenz VA Medical Center (CMC-VAMC, ACORP: 02004) and the University of Texas at Austin (UTA; Protocol # AUP20252200278, approved 02/07/2025) Institutional Animal Care and Use Committee and conducted according to the NIH Guide for the Care and Use of Laboratory Animals. Adult male Sprague-Dawley rats (300–330 g, Charles River Laboratories) were used at UPenn for *ex vivo* experiments. Adult male (300-500 g) and female (200-250 g) Sprague-Dawley rats were used at UTA for *ex vivo* experiments because previous studies observed no sex differences in functional and morphological outcomes in *ex vivo* stored nerves^61^.

#### Sciatic nerve harvest and culture

Rats were euthanized through CO₂ asphyxiation followed by decapitation at UPenn or deep anesthesia with isoflurane followed by intracardiac injection of saturated potassium chloride at UTA. Sciatic nerves were swiftly dissected from the mid-thigh to the trifurcation under sterile conditions. Nerves were cut into 10 mm segments and transferred to 24-well culture plates at UPenn or cut into 18-23 mm segments and transferred to 6-well culture plates at UTA. Media were composed of DMEM (Thermo Fisher, Cat #11995065) supplemented with 10% fetal bovine serum (FBS), 100 U/mL penicillin, and 100 µg/mL streptomycin. Cultures were maintained at 37°C in a humidified atmosphere with 5% CO₂.

#### Boldine treatment

Boldine was obtained from Sigma-Aldrich (catalog #B3916, 98% purity by HPLC). Stock solutions (100 mM) were prepared in dimethyl sulfoxide (DMSO) and then diluted to working concentrations (10–100 μM) in culture medium (final DMSO concentration <0.1%). Vehicle control cultures received the same volume of DMSO. The media were replaced daily with fresh ones containing boldine or vehicle for the duration of the experiment (3 or 7 days).

#### Immunofluorescence staining

At the designated time points, nerve explants were fixed in 4% paraformaldehyde (PFA) in phosphate-buffered saline (PBS) for 1 hour at room temperature, cryoprotected in 30% sucrose overnight at 4°C, and embedded in OCT compound. Longitudinal cryosections (20 μm) were cut on a cryostat, mounted on Superfrost Plus slides, and stored at −80°C until use. For immunostaining, sections were permeabilized with 0.3% Triton X-100 in PBS for 10 minutes, blocked with 5% normal horse serum (NHS) for 1 hour, and incubated overnight at 4°C with chicken neurofilament-light conjugated to Alexa-Fluorophore (NF-L, 1:500). Sections were mounted with Fluoromount-G and imaged using a confocal microscope. Fluorescence thresholds were set using the Otsu method in ImageJ and applied uniformly across all images.

#### Axon density quantification

Axon density was quantified by performing analyses on four randomly selected regions of interest (ROIs) per explant (n = 4 explants per group)^61^. Quantification was performed blinded to treatment groups.

#### Morphological analysis

After 7 days of storage, nerve explants were retrieved from culture plates, washed several times in saline, and assessed for gross morphology. Sample length after storage was measured and expressed as a ratio relative to length immediately after harvest. Nerve explants were also assessed for curling based on gross appearance.

Immediately following gross morphological assessment, 4-5 mm segments from the middle of the nerve explants were harvested. Samples were fixed in 2% PFA/3% glutaraldehyde (Electron Microscopy Sciences [EMS], Cat #15713, 16220) in 0.1 M sodium cacodylate buffer (EMS, Cat #11653) and treated and embedded.^61^ Semi-thin sections (0.5 μm) were cut with glass knives on a UC7 ultramicrotome (Leica), stained with toluidine blue, and imaged on an SP8 confocal microscope (Leica).

Axons were classified into five morphological categories and reported as a percentage of total axon count using ImageJ.^62^ Analysis was performed on four randomly selected ROIs per explant (n = 3 explants per group), with >750 axons quantified per group.

#### Statistical analyses

All statistical analyses were performed using GraphPad Prism 10 (GraphPad Software). Axon densities were compared using one-way ANOVA with Tukey’s post-hoc test. Normalized lengths of stored nerve explants were compared using Welch’s t-test. Percentages of curled nerve explants were compared using Chi-Square test. Morphological analysis of axons was performed using two-way ANOVA with Sidak’s post-hoc test. p < 0.05 was considered statistically significant. Data are presented as mean ± SEM or percentages as indicated in respective figure legends.

## Results

### Boldine Inhibits SARM1 NADase Activity in a Fluorescence Polarization Assay

Direct inhibition of SARM1 enzyme activity was measured by quantifying NAD⁺ hydrolysis using the Transcreener ADPR fluorescence polarization (FP) assay, which tracks ADPR buildup during SARM1-mediated NAD⁺ cleavage.

Enzyme titration experiments were initially performed to identify assay conditions that kept the reaction within the linear initial-velocity range (<20% substrate conversion). Increasing SARM1 concentrations caused a strong, dose-dependent change in fluorescence polarization signal, with an EC₅₀ of approximately 2.5 nM, indicating good assay sensitivity and dynamic range (**Fig. 2A**). An ADPR calibration curve, showing a gradual decrease in polarization with increasing ADPR levels, enabled quantitative conversion of FP signals to the amount of product formed (**Fig. 2B**). Under optimized conditions, dose-response analysis of boldine against 40 nM SARM1 revealed concentration-dependent inhibition with an IC₅₀ of about 7.5 μM (**Fig. 2C**). These findings demonstrate that boldine can inhibit SARM1’s catalytic activity in vitro, although the precise inhibitory mechanism needs further investigation.

**Fig. 2.**
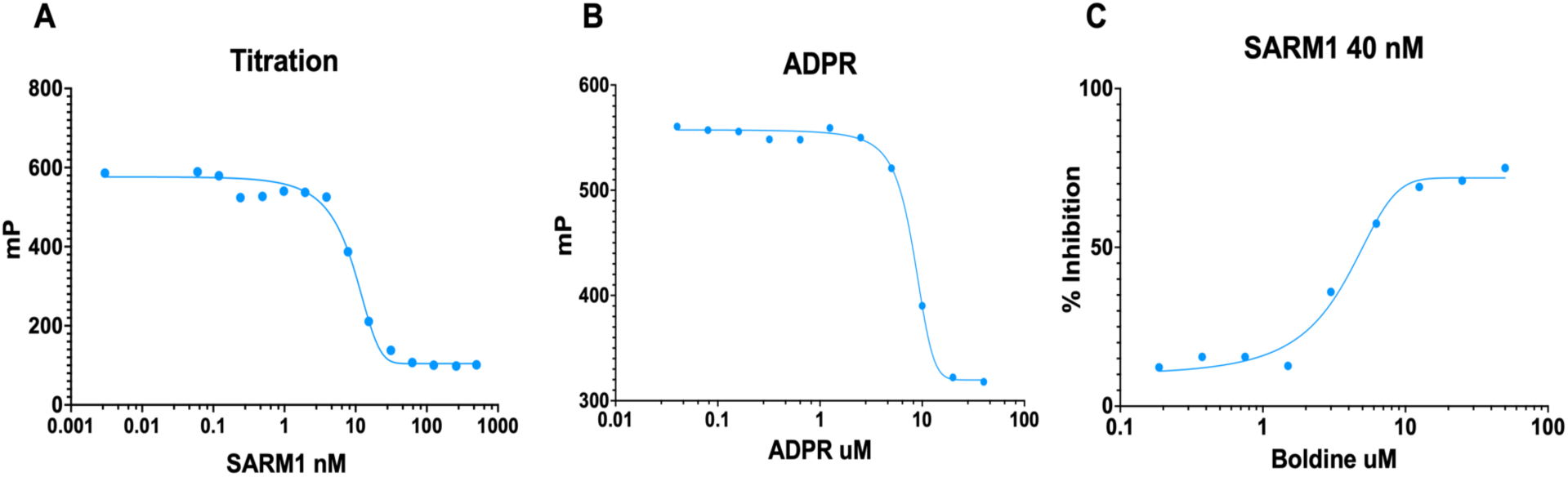
SARM1 Activity and Boldine Inhibition Using the Transcreener ADPR FP Assay. (A) SARM1 enzyme titration measured with the Transcreener ADPR fluorescence polarization (FP) assay. Increasing enzyme concentrations produced a dose-dependent change in the FP signal, with an EC₅₀ of about 2.5 nM. Titration experiments helped identify assay conditions that maintained reactions within the initial-velocity range (<20% substrate conversion). (B) ADPR standard curve used to convert FP signals into product concentration, allowing estimation of SARM1-mediated NAD⁺ hydrolysis. (C) Dose–response analysis of boldine inhibition of SARM1 NADase activity using 40 nM SARM1. Nonlinear regression analysis estimated an IC₅₀ of around 7.5 μM; each assay condition was performed in duplicate and data shown are means of the duplicates.

### Structural Organization of SARM1 in Active and Inactive Conformations

Cryo-electron microscopy structures of SARM1^16, 19^, representing inactive and active conformational states, were analyzed to identify a structural basis for ligand binding. Models were visualized using UCSF Chimera.^37^

In the autoinhibited structure (PDB ID: 7CM6), SARM1 forms an octameric ring in which the peripheral ARM regulatory domains surround and spatially restrict the central TIR NADase domains, consistent with the suppression of the catalytically active interface (**Fig. 3A**). Side-view projections illustrate the layered organization of ARM, SAM, and TIR domains within this assembly (**Fig. 3B**). The ARM regulatory region shows multiple surface-accessible pockets that could accommodate small-molecule ligands capable of altering their conformation (**Fig. 3C**). Overall, these structures aid in identifying potential binding sites on SARM1.

**Fig. 3.**
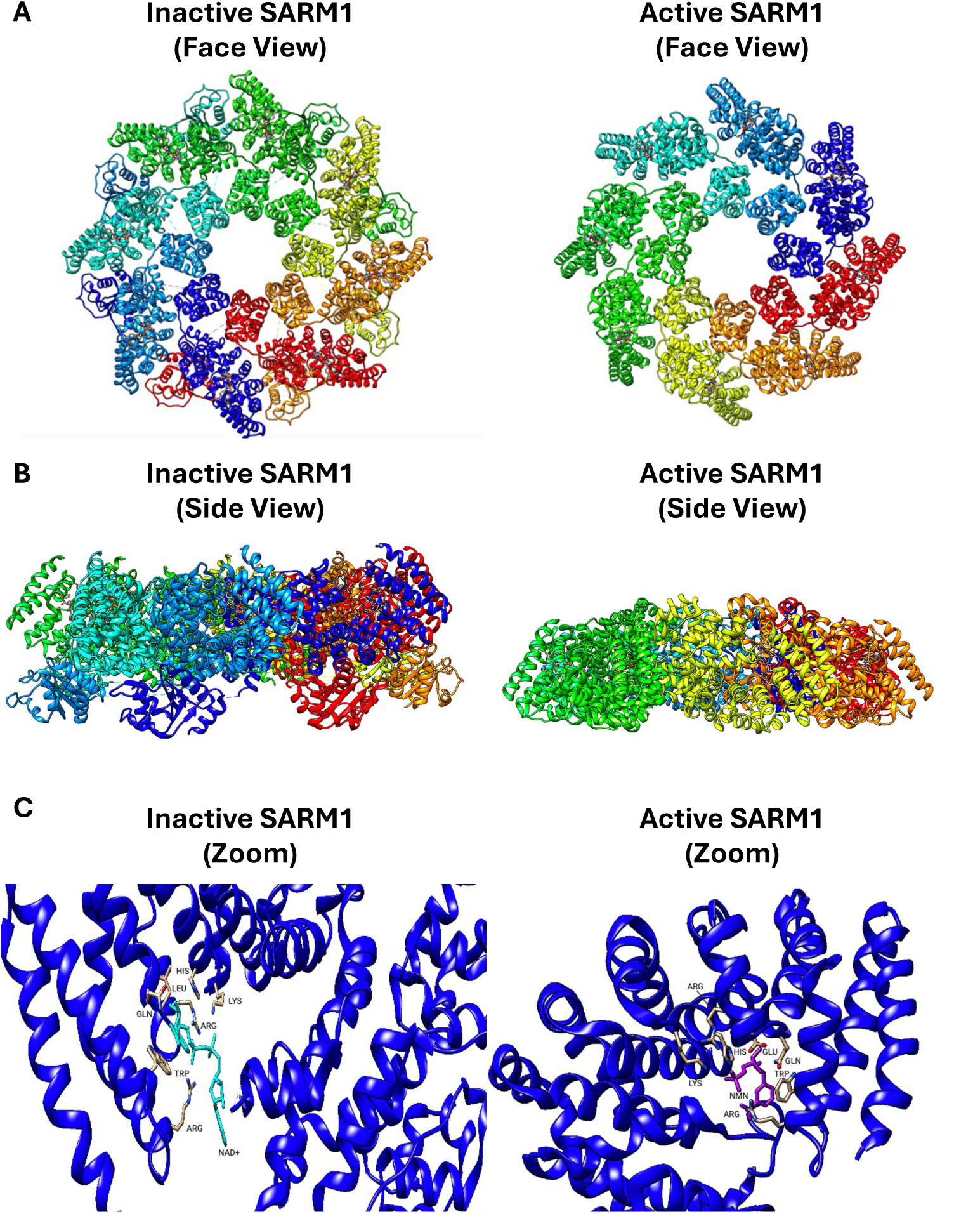
Structural States of SARM1. Structural models of SARM1 visualized with UCSF Chimera. (A) Top-view and (B) side-view of the homo-octameric assembly of the autoinhibited and active SARM1 complex (PDB ID: 7CM6 and 7NAL), as characterized by cryo-electron microscopy. Individual protomers are colored to show quaternary organization and stacked architecture of the ARM regulatory, SAM oligomerization, and TIR catalytic domains. (C) Surface representation of the ARM regulatory region displaying surface-accessible pockets identified by computational analysis as candidate small-molecule binding sites; shown here based on our own structural modeling using the PDB structures indicated. Pockets were visualized to identify regions suitable for targeted docking of boldine and reference compounds.

### Docking Simulations Identify Potential Binding Sites Across Multiple SARM1 Conformations

Molecular docking simulations were conducted using GNINA^33, 40^ across several experimentally determined SARM1 structures that represent different conformational and domain states, including the autoinhibited full-length structure (PDB ID: 7CM6), the activated full-length structure (PDB ID: 7NAL), the isolated ARM regulatory domain (PDB ID: 7M6K), and the catalytic TIR NADase domain (PDB ID: 6O0Q).^7, 9, 13, 14, 16, 19, 35, 60^

When docked against the autoinhibited structure (PDB ID: 7CM6), boldine was predicted to adopt a favorable position within a pocket near the ARM regulatory domain (**Fig. 4A**). Among all tested compounds, including the reference inhibitor DSRM-3716, boldine showed the strongest predicted binding affinity (**Fig. 4B**). This pose had a predicted binding free energy of ΔG ≈ −8.3 kcal/mol, corresponding to a K*d* of about 0.76 μM.

**Fig. 4.**
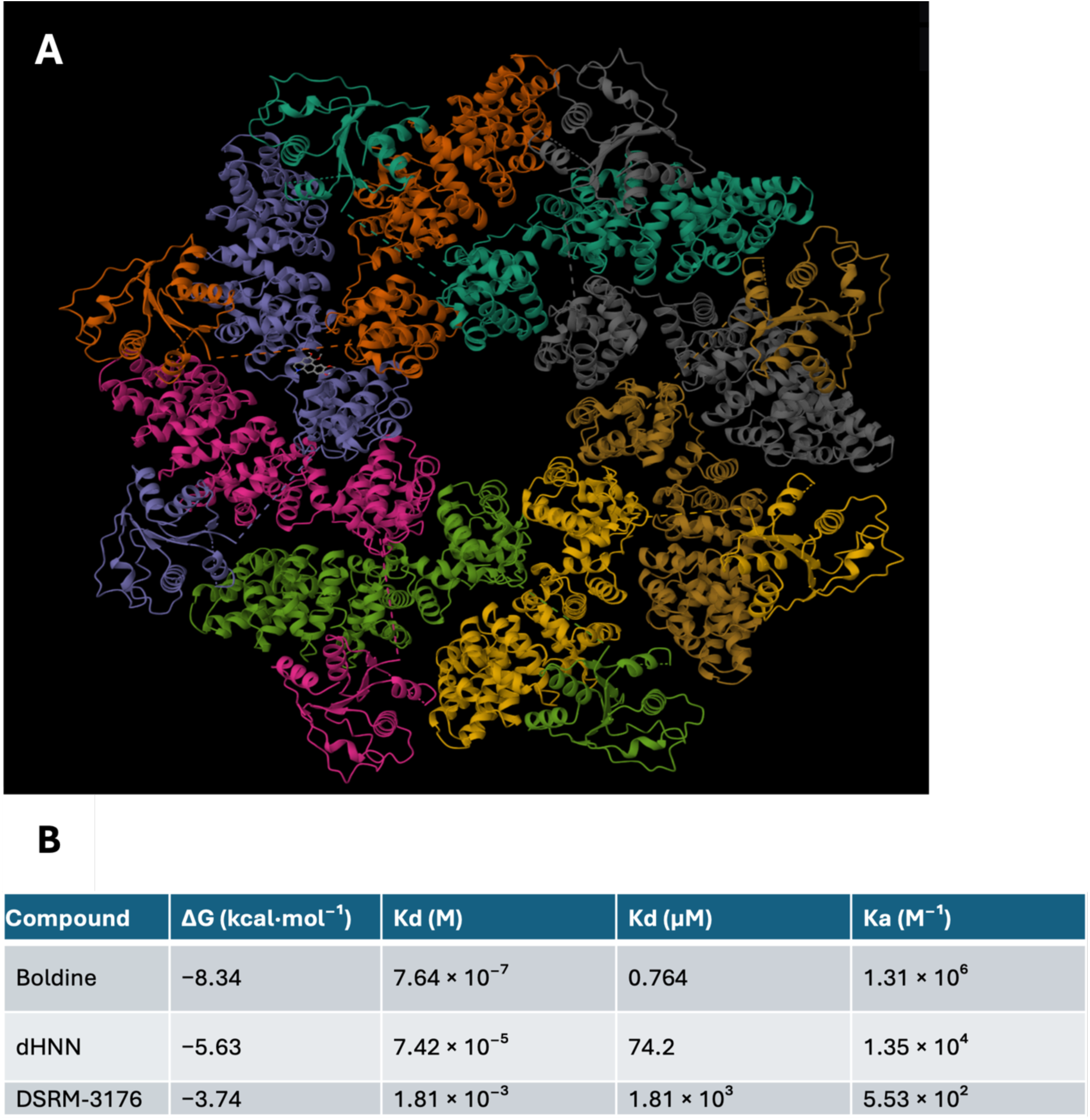
Docking Predictions for Boldine Binding to Inactive SARM1. (A) Predicted docking pose of boldine within the autoinhibited SARM1 structure (PDB ID: 7CM6). (B) Comparative docking metrics for boldine and reference compounds. Predicted binding affinities (Kᵈ) were estimated from calculated free energies (ΔG) obtained from docking simulations. Docking procedures and scoring functions are described in the Methods.

Docking against the activated full-length structure (PDB ID: 7NAL), in which ARM domain displacement is thought to enable TIR oligomerization into a NAD⁺-hydrolyzing state, identified ligand-accessible pockets (**Fig. 5A**), although predicted affinities were lower than in the autoinhibited state. The top-scoring pose resulted in a ΔG of approximately −6.6 kcal/mol (K*d* around 13 μM) (**Fig. 5B**).

**Fig. 5.**
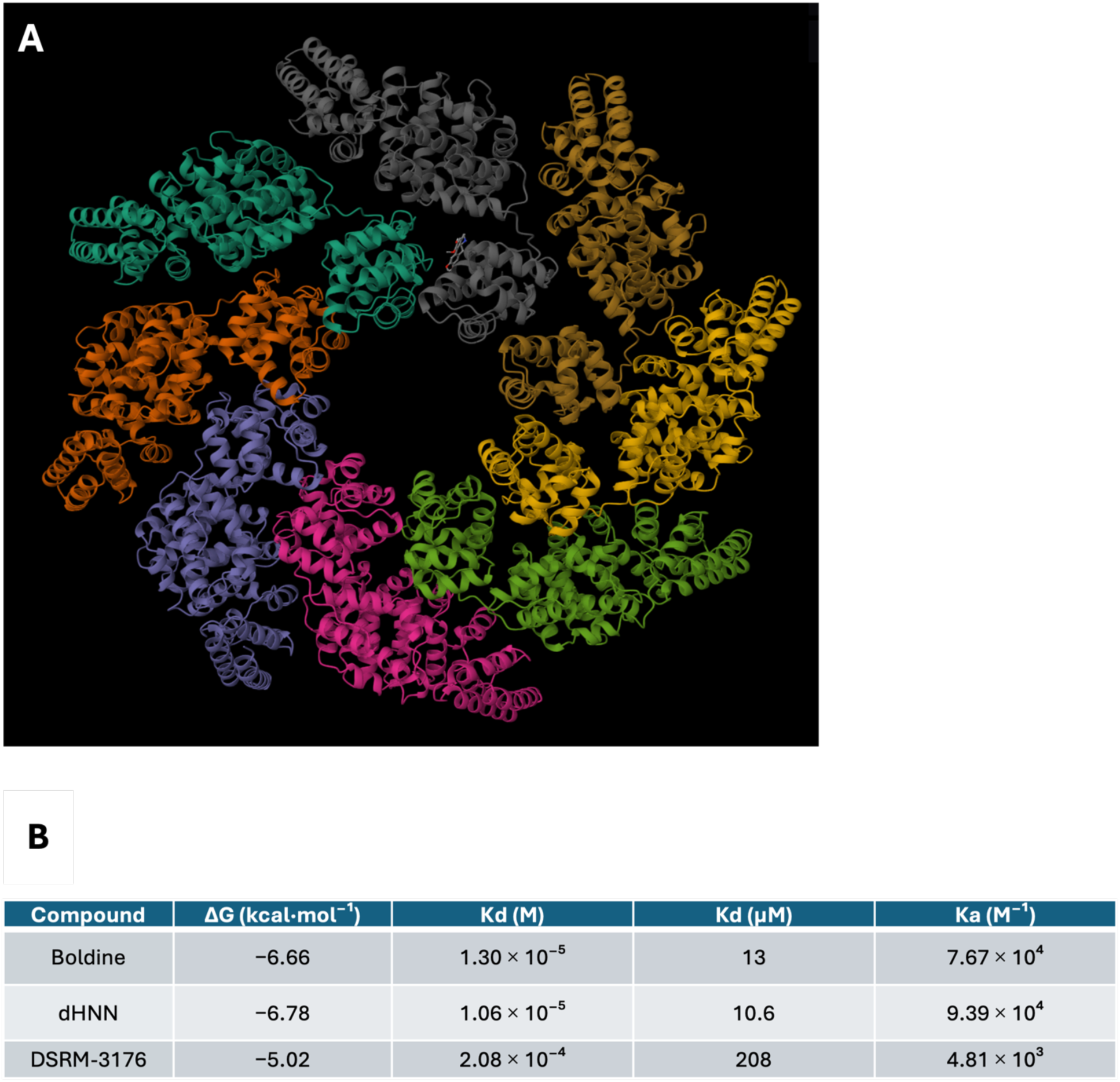
Docking Predictions for Boldine Binding to Activated SARM1. (A) Predicted docking pose of boldine within the activated full-length SARM1 structure (PDB ID: 7NAL). (B) Comparative docking metrics for boldine and reference compounds derived from calculated ΔG values.

Simulations against the isolated ARM regulatory domain (PDB ID: 7M6K) positioned boldine in a surface-accessible pocket consistent with regulatory ligand binding (**Fig. 6A**). The top configuration yielded ΔG ≈ −6.7 kcal/mol (K*d* ≈ 12 μM), with boldine again surpassing DSRM-3716 (**Fig. 6B**).

**Fig. 6.**
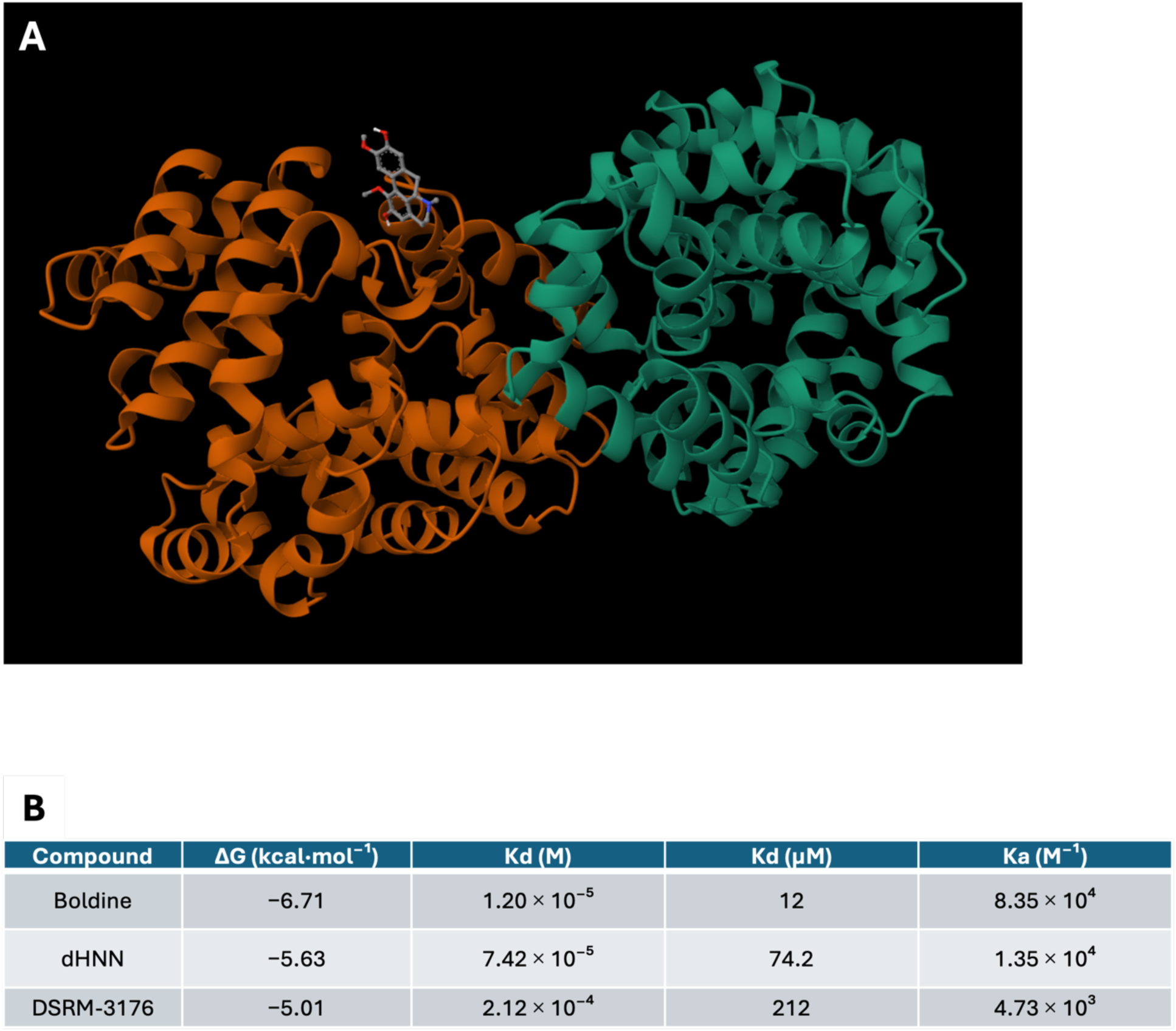
Predicted Binding of Boldine to the SARM1 ARM Regulatory Domain. (A) Docking pose of boldine within the isolated ARM regulatory domain structure (PDB ID: 7M6K). (B) Comparative docking metrics for boldine and reference compounds, with predicted dissociation constants derived from calculated ΔG values.

Against the isolated catalytic TIR domain (PDB ID: 6O0Q), multiple energetically favorable poses were identified near the catalytic region (**Fig. 7A**). The highest-scoring pose yielded ΔG ≈ −6.6 kcal/mol (K*d* ≈ 13.5 μM); comparative metrics (**Fig. 7B**) again favored boldine over the reference compound.

**Fig. 7.**
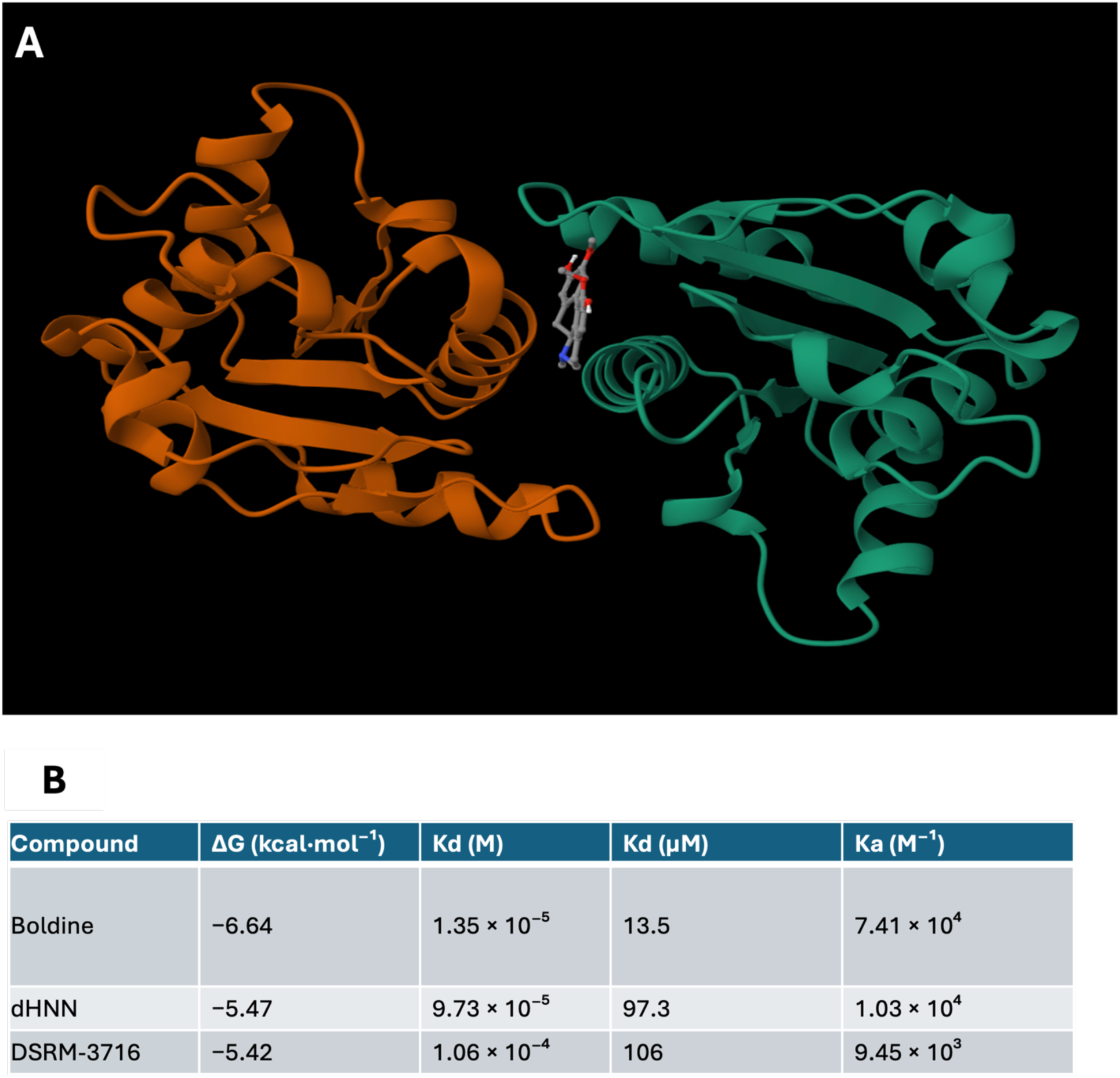
Predicted Binding of Boldine to the Catalytic TIR Domain of SARM1. (A) Predicted docking pose of boldine within the catalytic TIR domain (PDB ID: 6O0Q). (B) Comparative docking metrics for boldine and reference compounds derived from docking simulations.

The predicted pose places boldine near catalytically relevant residues (Glu642, His685, and Asn640), consistent with partial steric occlusion of the NAD⁺ binding cleft and/or conformational perturbation of the catalytic site, rather than exclusive reliance on passive steric exclusion alone. The experimentally measured IC₅₀ (≈ 7.5 μM) falls within one order of magnitude of the docking-derived affinities (K*d* ≈ 0.76–13 μM), providing partial support for the predicted binding mode, though it is not sufficient on its own to confirm it.

### Boldine Preserves Axonal Integrity in *Ex vivo* Culture

The potential of boldine to prevent axonal degeneration in intact tissue was tested using a well-established *ex vivo* Wallerian degeneration model. Transected rat sciatic nerve segments were cultured with daily medium changes and treated with either boldine from two separate sources or a vehicle (**Fig. 8A**).

**Fig. 8.**
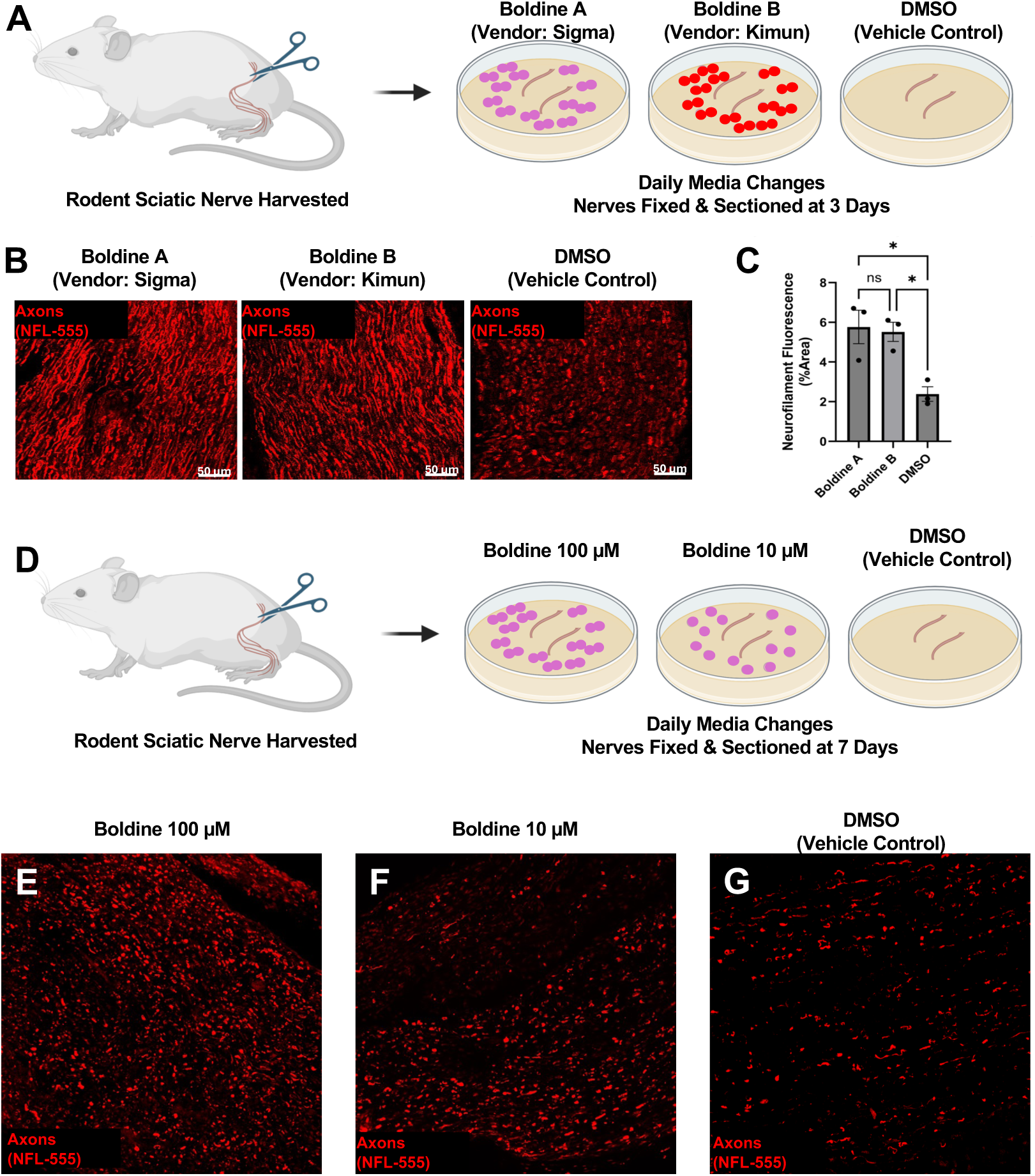
Boldine Preserves Axonal Integrity in an *ex vivo* Sciatic Nerve Explant Model. (A) Schematic of the rat sciatic nerve explant model. Sciatic nerves were harvested, cultured *ex vivo* with daily media changes, and fixed 3 days after transection. Experimental treatment groups include vehicle control (DMSO) and boldine from two independent vendors. (B) Representative immunofluorescence images of axons labeled with neurofilament light chain (NFL-555) at 3 days post-explant. Scale bar: 50 μm. (C) Quantification of neurofilament-positive area normalized to total tissue area. Data are shown as mean ± SEM (n = 3 explants per group). Both boldine preparations led to significantly greater neurofilament-positive area compared to vehicle control, with no significant difference between boldine sources. *p<0.05, Tukey’s multiple comparisons test. (D) Experimental schematic of sciatic nerve explants cultured for 7 days after transection and treated with boldine (10 μM or 100 μM) or vehicle control, with daily media replacement. (E-G) Representative images of NFL-555–labeled axons at Day 7. Boldine-treated explants appeared to maintain more continuous neurofilament labeling than DMSO-treated controls, which showed fragmented axonal structures indicative of progressive degeneration.

Neurofilament light chain immunolabeling (NFL-555) served as the primary indicator of axonal integrity. By day 3, nerves treated with vehicle showed significant fragmentation and disorganized neurofilament structure, while boldine-treated explants preserved much more continuous labeling across the nerve bundles (**Fig. 8B**).

Quantification of the neurofilament-positive area, normalized to the total tissue cross-section, confirmed greater axonal preservation in boldine-treated explants (**Fig. 8C**). Vehicle-treated nerves achieved only 2.38 ± 0.37% neurofilament-positive area (mean ± SEM; n = 3); Boldine A (Sigma) increased this to 5.77 ± 0.84%, and Boldine B (Kimun) to 5.52 ± 0.49% (n = 3 explants per group). Both preparations differed significantly from the vehicle (Tukey’s post-hoc tests; Boldine A vs. DMSO, p = 0.017; Boldine B vs. DMSO, p = 0.0237), and the two sources were statistically indistinguishable (Tukey’s post-hoc test, p = 0.9539), supporting consistent bioactivity regardless of origin.

To assess whether protection persists over time, explants were maintained for 7 days with boldine (10 μM or 100 μM) or vehicle, with daily medium changes (**Fig. 8D)**. On day 7, neurofilament imaging showed nearly complete loss of axonal structure in vehicle-treated explants, whereas nerves treated with boldine retained organized axonal morphology (**Fig. 8E-8G**). Neurofilament-positive area at Day 7 was quantified using the same blinded Otsu thresholding approach applied to Day 3 data (n = 3 explants per group). These results suggest that boldine may protect axonal structure in an *ex vivo* sciatic nerve degeneration model, and that this protective effect persists for at least 7 days after injury, maintaining neurofilament immunoreactivity and axon continuity. These findings are consistent with the in vitro enzymatic inhibition of SARM1 and support the hypothesis that boldine could suppress pathways involved in axonal degeneration.

### Boldine reduces gross and axonal disintegration in *ex vivo* cultures

To assess axonal morphology in ex vivo culture, we maintained 16 additional transected rat sciatic nerve explants at 37°C for 7 days in medium containing boldine (100 μM) or vehicle (0.1% DMSO), with daily medium changes. On Day 7, before fixation and embedding, we observed gross differences between treatment groups (**Fig. 9A**). Interestingly, 4 of 8 vehicle control explants became curled, whereas none of the 8 boldine-treated explants curled (Chi-square test, p< 0.05, **Fig. 9B**). Explants in both groups were also shorter on Day 7 than immediately after harvest. However, boldine-treated explants retained a significantly greater proportion of their initial length than vehicle-treated explants (0.799 ± 0.014 vs. 0.700 ± 0.025, mean ± SEM; Welch’s t-test, p = 0.0059) (**Fig. 9C**).

**Fig. 9.**
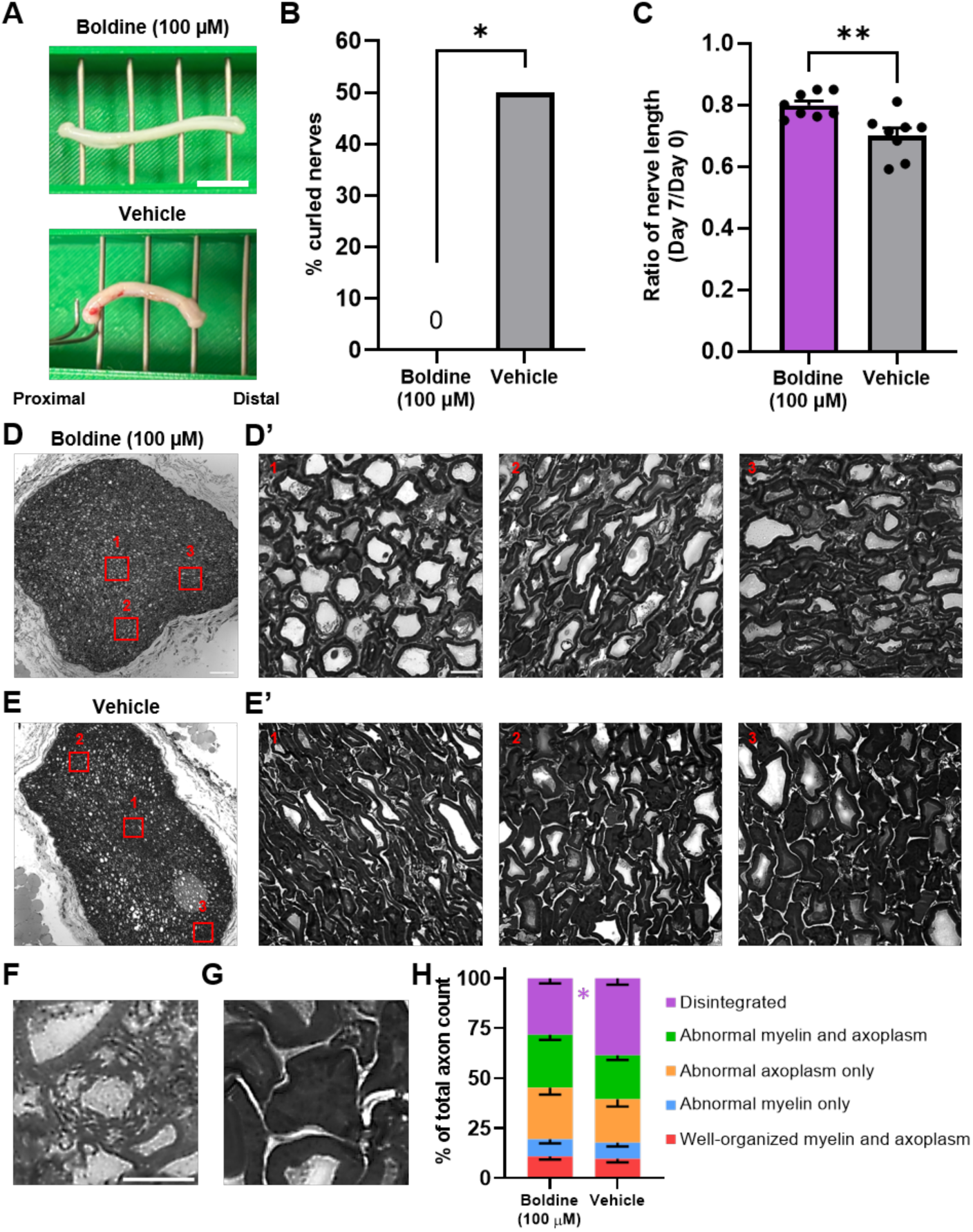
Boldine treatment preserves nerve explant structures. (A) Representative images of stored nerve explants after 7 days of culture with 100 µM boldine (vendor: Sigma) or vehicle control (0.1% DMSO) with daily media replacement. Scale bar = 5 mm. n = 8 nerves per group. (B) Percent of stored nerve explants that became curled, as illustrated in vehicle control in panel A. *p<0.05, Chi-Square test. (C) Ratio of nerve explant length normalized to their length prior to storage. **p<0.01, Welch’s t-test. (D-E) Representative low-magnification images of semi-thin sections (0.5 µm) of nerve explants at Day 7. Scale bar = 100 µm. (D’-E’) High-magnification insets of the boxed regions in D-E. Scale bar = 10 µm. (F-G) Representative images of different types of disintegrated axons. (F) Fragmented axon. (G) Collapsed axon. Scale bar = 5 µm. (H) Percentages of axons according to different categories defined by Zhou et al.^62^. *p<0.05, Sidak’s multiple comparisons test. n = 12 ROIs per group (4 ROIs x 3 samples), with >750 axons quantified per group.

We next examined whether these gross differences were accompanied by differences in axonal morphology. After 7 days of culture, we collected 4–5-mm segments from the middle of each nerve explant and prepared semi-thin toluidine blue–stained transverse sections. Figures 9D and 9E show representative low-magnification sections from a boldine-treated explant and a vehicle control explant, respectively.^62^ Both treatment groups contained a range of well-organized, abnormal, and disintegrated axonal profiles (**Fig. 9D′–E′)**.

To illustrate the heterogeneity observed within each sample, panels 9D′–E′ show three high-magnification regions from the representative explants. Within each condition (Boldine or Vehicle), region 1 shows relatively better-preserved morphology, region 2 shows more abnormal profiles, and region 3 shows the most severe degenerative changes. The most severely affected regions contain numerous disintegrated axons. Figures 9F and 9G show two morphologies of axonal disintegration: fragmentation and collapse, respectively. In the collapsed axon, the myelin sheath has invaginated to occupy the axoplasm completely.

Blinded morphological classification of 800 axons in the boldine group and 750 axons in the vehicle group (1,550 axons total) showed that boldine treatment significantly reduced the proportion of disintegrated axons relative to vehicle treatment (28.18 ± 2.56% vs. 38.58 ± 3.17%, mean ± SEM; Sidak’s post-hoc test, p =0.0269) (**Fig. 9H**). Collectively, these findings show that daily treatment with 100 µM boldine prevented nerve curling, reduced gross nerve shortening, and impeded axonal disintegration after 7 days of ex vivo culture.

## Discussion

Across computational, biochemical, and *ex vivo* experiments, boldine treatment was associated with inhibition of SARM1 NADase activity and preservation of axonal integrity following nerve injury. In a rat sciatic nerve explant model, boldine maintained axonal structure at both 3 and 7 days after transection, consistent with suppression of degeneration pathways that normally drive rapid axonal breakdown. These findings, together with biochemical evidence of SARM1 inhibition, support a model in which boldine delays the initiation of programmed axonal degeneration.^5, 8, 42^

### Mechanistic Interpretation of Dual-Site SARM1 Inhibition

Computational modeling suggests that boldine may engage SARM1 at both the TIR catalytic site and the ARM–TIR regulatory interface, raising the possibility of combined competitive and allosteric inhibition.^14, 43, 44^ While direct structural validation is required, this dual-site interaction could provide a mechanistic basis for sustained inhibition across activation states of the enzyme. If confirmed, such a mechanism would distinguish boldine from existing inhibitor classes and offer a strategy for more effective suppression of axonal degeneration.

### Comparison to Existing SARM1 Inhibitor Classes

SARM1 inhibitor classes differ in their targets and limitations. Base-exchange inhibitors like DSRM-3716 (5-iodoisoquinoline) are converted by SARM1 itself into a potent orthosteric inhibitor (1AD) through a base-exchange reaction with NAD⁺. While this class is potent and selective, DSRM-3716 and related compounds have been shown to paradoxically activate SARM1 at sub-inhibitory concentrations, accelerating neurite degeneration and potentially narrowing the therapeutic window.^41^ Covalent inhibitors targeting ARM-domain cysteines (notably Cys311) offer long-lasting suppression in preclinical settings, but concerns about selectivity and potential immune responses from haptenized proteins remain unresolved.^25, 45^ Base-exchange inhibitors are innovative mechanistically, but their tendency to paradoxically activate SARM1 at low exposure levels raises concerns in situations where continuous target coverage is not feasible.^26^ Critically, a recent study demonstrated that known base-exchange inhibitors, including DSRM-3716, paradoxically activated SARM1 at sub-inhibitory concentrations, accelerating neurite degeneration. These findings underscore the need for alternative scaffolds, such as boldine, that engage SARM1 through distinct mechanisms without this pro-degenerative risk.^59^ Boldine lacks reactive functional groups expected to covalently modify SARM1, making non-covalent inhibition the most plausible mechanism. However, formal reversibility has not yet been demonstrated and will require kinetic experiments varying substrate concentration at fixed boldine concentrations. An estimated selectivity for the autoinhibited over the active enzyme suggests the autoinhibited version stabilizes the resting state without disrupting NAD⁺ homeostasis in healthy neurons^16, 19^. However, this hypothesis needs to be tested in intact cellular systems given NAD⁺’s essential metabolic roles.

### Reconciling Boldine’s Established Neuroprotective Mechanisms

Prior studies of boldine have largely attributed its neuroprotective effects to modulation of connexin hemichannels, inflammation, and oxidative stress.^29, 30^ In models of delayed repair following peripheral nerve injuries, boldine reduces connexin 43/45 expression, attenuates muscle atrophy, and enhances electrophysiological recovery following delayed neurorrhaphy. ^59^ In spinal cord injury models, boldine treatment has been shown to increase spared white matter and improve locomotor recovery, accompanied by reduced glial activation and increased expression of genes associated with axonal growth and synaptic function.^47, 48^ Similarly, in ALS models, boldine reduces connexin hemichannel activity and oxidative stress while improving motor neuron survival and locomotor performance.^49^ Collectively, these findings demonstrate consistent neuroprotective effects across injury and disease contexts but have been interpreted primarily as downstream actions at the level of muscle, glia, and inflammatory signaling.

However, connexin hemichannel activity is increasingly recognized as a contributor to denervation-induced muscle atrophy and impaired reinnervation, linking these pathways more directly to axonal stability.^50^ In this context, the current findings raise the possibility that previously reported effects of boldine may reflect, at least in part, upstream preservation of axons through modulation of degeneration pathways. Specifically, quantal release of ACh from terminal boutons suppresses the *de novo* appearance of Cx43/Cx45 hemichannels in the sarcolemma. When nerve injury eliminates this ACh release, hemichannels emerge and initiate the signaling cascades driving denervation-induced muscle atrophy, an effect attenuated by double Cx43/Cx45 conditional knockout in myofibers or by the hemichannel blocker D4. What is mechanistically novel in the present work is the proposal that boldine acts upstream of this pathway by inhibiting SARM1 and thereby preventing the Wallerian degeneration that causes loss of ACh release in the first place. The identification of SARM1 inhibition as a potential mechanism provides a unifying framework connecting these observations. Preservation of axonal structure may maintain neuromuscular signaling, limit secondary inflammatory responses, and sustain tissue receptivity to reinnervation, thereby enabling the downstream benefits observed across models.

### Translational Considerations and Pharmacokinetics

Confirmation of the proposed dual-site mechanism would establish a clear molecular basis for using a plant-derived natural product in neuroprotection. Boldine has a long history in traditional South American medicine, particularly in boldo-leaf preparations, and preclinical toxicology data generally support a tolerable safety profile, although comprehensive dose–response and long-term toxicity studies remain limited.^29, 30^ The aporphine scaffold is synthetically accessible and chemically adaptable, providing a foundation for structure–activity relationship–guided optimization of potency, selectivity, metabolic stability, and tissue targeting.^31, 51^

Pharmacokinetic studies indicate that boldine is rapidly absorbed following oral administration, reaching peak plasma concentrations within approximately 30 minutes and accumulating in tissues, with highest levels in the liver and lower but measurable concentrations in the brain and heart.^30^ However, boldine undergoes rapid metabolism, with a reported plasma half-life in rats of approximately 30 minutes and relatively low oral bioavailability, likely due to significant first-pass hepatic metabolism. These properties suggest that, although boldine can cross the blood-brain barrier and affect CNS functions, the duration of CNS effects may be limited, particularly for conditions requiring sustained inhibition such as suppression of SARM1-mediated axon degeneration. Consequently, effective therapeutic application may require repeated dosing, long-acting formulations, or structural optimization to improve metabolic stability and tissue retention. Moreover, while boldine has demonstrated neuroprotective effects across multiple *in vivo* models, its ability to directly modulate canonical axon degeneration pathways, such as SARM1-mediated NAD⁺ depletion, has not yet been established.

### Limitations

The current study has several important limitations. While *in silico* predictions are well supported by RMSD values compared to known inhibitors, *in silico* predictions cannot replace cryo-EM or crystallographic structures of boldine–SARM1 complexes; such data are necessary to confirm dual-site binding directly. The biochemical IC₅₀ of 7.5 μM indicates activity against recombinant SARM1 but may differ from how the enzyme behaves in intact neurons. Additionally, the separate roles of the two proposed binding sites in neuroprotection must be tested experimentally using ΔARM and catalytic mutant (E642A/H685A) SARM1 constructs, along with competition assays that independently vary [NAD⁺] and [NMN].

Although boldine-treated explants exhibited improved gross morphology and reduced axonal disintegration after 7 days of *ex v*ivo culture, these results should be interpreted primarily as evidence of structural axon preservation rather than definitive preservation of electrophysiological function. The explant paradigm used here exposed transected nerves to prolonged culture at near-physiological temperature in DMEM-based medium, conditions that model degenerative stress but do not reproduce the vascular, metabolic, and cellular support present *in vivo*. Nor do these conditions reflect optimized peripheral nerve storage protocols. Indeed, prior work has demonstrated that rat peripheral nerve segments/allografts stored at 4°C in optimized diluted physiological solutions can retain compound action potential conduction and axonal morphology for extended periods^61–62^.Therefore, the current findings support a protective effect of boldine on nerve structure under stringent ex vivo conditions, while the relationship between this structural preservation and recoverable electrophysiological conduction will require follow-up studies in either optimized storage paradigms or in vivo injury models. An additional limitation is that these explant studies were not designed or powered to assess sex as a biological variable. Accordingly, potential sex-dependent differences in boldine-mediated axon preservation remain unresolved and should be evaluated in future in vivo and ex vivo studies with appropriately balanced cohorts.

An additional limitation is that, while structural preservation of axons was demonstrated, functional electrophysiological activity was not directly assessed, leaving the relationship between structure and signal transmission unresolved. Notably, prior findings in a rodent *in* vivo denervation model showed that oral boldine treatment initiated immediately after nerve transection preserved evoked muscle responses at 2 and 6 weeks post-injury, suggesting maintained neuromuscular connectivity.^46^ However, direct coupling of structural and electrophysiological outcomes within the same experimental system will be required to definitively establish this relationship.

### Future Directions

Future priority areas include SAR-guided analog optimization to improve potency and metabolic stability; IND-enabling toxicology focused on potential pro-oxidant effects at supratherapeutic doses; NAD⁺ metabolite profiling (NAD⁺, NMN, ADPR, cADPR) in vitro and to confirm on-target SARM1 engagement in vivo.

### Conclusion

This study identifies boldine as a potential modulator of programmed axonal degeneration, supported by computational, biochemical, and *ex vivo* evidence demonstrating inhibition of SARM1 NADase activity and preservation of axonal structure following injury. These findings extend the current understanding of boldine’s neuroprotective effects beyond modulation of connexin hemichannels, inflammation, and oxidative stress, and suggest a potential upstream role in delaying the initiation of axon degeneration. Computational modeling raises the possibility of dual-site engagement of SARM1 across both catalytic and regulatory domains, providing a mechanistic framework for sustained inhibition across activation states; given that SARM1 functions as a homo-octamer, binding of boldine at either site on one protomer may additionally propagate allosteric effects across the multimeric complex. While direct structural validation and in vivo confirmation of SARM1 pathway modulation remain necessary, and the relationship between structural preservation of axonal integrity and functional recovery requires further investigation, this work positions boldine as a promising candidate for pharmacologic strategies aimed at preserving axonal integrity and extending the therapeutic window following nerve injury.

## Author Contributions

J.C.B. and T.T.L. contributed equally to this work. J.C.B. and D.K.C. conceived the study. T.T.L. performed computational experiments, including molecular docking and structural modeling. J.C.B., H.G., D.R.C., M.M.T., L.Z., C.A.T., A.N.O., P.N., and C.Z.Y. performed experiments. J.C.B., T.T.L., H.G., and L.Z. performed data analysis and figure preparation. J.C.B. and T.T.L. wrote the manuscript with input from all authors. All authors reviewed and edited the manuscript. D.K.C., C.P.C., and G.D.B. supervised the project and provided funding support.

## Data Availability

Raw data are available from the corresponding author upon reasonable request. All data supporting the findings of this study are available within the article. Structural data used for docking were obtained from the RCSB Protein Data Bank (www.rcsb.org) under accession codes 7CM6, 7NAL, 7M6K, and 6O0Q.

## Acknowledgements

Funding was provided by the Department of Veterans Affairs [Merit Review I01-RX005045 and Center Grant I50-RX004845] and the National Institutes of Health [NIAMS R01-AR083489] to D.K.C and. [NINDS R01-NS128086] to G.D.B. We thank the Corporal Michael J. Crescenz VA Medical Center and James J. Peters VA Medical Center for their support. The opinions, interpretations, conclusions, and recommendations expressed herein are those of the authors and are not necessarily endorsed by the Department of Veterans Affairs or the National Institutes of Health.

## Ethics Statement

The animal study was approved by the Corporal Michael J. Crescenz VA Medical Center and University of Texas at Austin Institutional Animal Care and Use Committees (IACUC).

